# A cellular assay system revealed the deamidase-mediated regulation of the Arg-mediated N-end rule protein degradation pathway

**DOI:** 10.1101/2024.03.11.584350

**Authors:** Shusei Yoshida, Ren Ohta, Mami Miyabe, Taku Tamura

## Abstract

Regulated protein production and degradation are essential for maintaining proteostasis in eukaryotic cells. The N-end rule, or N-degron pathway, is a protein degradation machinery in which the N-terminal amino acid is the mark of the protein degradation via the proteasome. The N-end rule is a conserved protein disposal machinery in eukaryotic cells. However, the precise cellular mechanisms and their physiological roles are not fully understood. Herein, we report the development of an Arg-mediated N-end rule assay system using artificial substrates expressed in cultured cell lines. We demonstrated that the N-end rule degradation is significantly influenced by the expression levels of N-terminal amino acid-modifying enzymes, including NTAN1, NTAQ1, and ATE1. In the N-terminal Asn protein pathway, an increase in NTAN1 or ATE1 expression promotes its disposal via the N-end rule degradation pathway. Interestingly, overexpression of NTAQ1 decreased the degradation of the protein bearing Gln at the N-terminus. Computational prediction of NTAQ1 and ATE1 complex formation suggests that the outer loop region of NTAQ1 is involved in its interaction with ATE1 and that the NTAQ1 overexpression may negatively affect this interaction. Our findings suggest that the degradation activity of the Arg/N-end rule pathway is positively or negatively regulated by deamidase expression levels and propose a higher degree of control of protein degradation by the Arg/N-end rule within cells.

## Introduction

Protein degradation is important for maintaining the balance of protein quality and quantity within cells. A variety of proteolytic mechanisms are executed cooperatively in eukaryotic cells. Selective and regulated degradation of undesired proteins, such as protein aggregates or overexpressed proteins, is important for the quality of life as it prevents the accumulation of protein aggregates, which is closely associated with conformational diseases, including neurodegenerative disorders (Balch et al., 2008). Proteasomes degrade polyubiquitinated proteins in various cellular locations, including the endoplasmic reticulum (ER), nucleus, mitochondria, and cytosol (Deshmukh et al., 2019). The polyubiquitination of target proteins is mediated by sequential post-translational modifications: E1 activates mono-ubiquitin, which is then recognized by E2, E3 captures substrate, and E2 conjugates mono-ubiquitin (Dikic & Schulman, 2023).

The N-end rule (or N-degron) is a protein-degradation mechanism that utilizes proteasomes. In the N-end rule pathway, the N-terminal amino acid of the target protein is a critical determinant for degradation (Varshavsky, 2019). Since the discovery of the N-end rule mechanism by Varshavsky et al. in 1985 (Bachmair et al., 1985), considerable progress has been made in elucidating the fundamental molecular mechanisms behind substrate formation, substrate recognition, and polyubiquitination machinery (Varshavsky, 2019).

The N-terminal Arg-mediated N-end rule branch (Arg/N-end rule) is a component of the eukaryotic N-end rule degradation pathway (Varshavsky, 2011). In the Arg/N-end rule, proteins bearing bulky hydrophobic or charged amino acid residues at the N-terminus are recognized as a substrate of the Arg/N-end rule (Varshavsky, 2011). The corresponding amino acids at the N-terminus of Arg/N-end rule substrates are generated by cellular proteases, followed by modifications in the case of Cys, Asn, Gln, Asp, or Glu (Figure 1A). E3s (Ubr1–7) recognize these specialized proteins and polyubiquitinate them for proteasomal degradation, in cooperation with E2s, Ube2a and Ube2b (Varshavsky, 2011). The functional roles of the N-end rule E3s and their molecular mechanisms have been well characterized in detail by structural analysis, and their importance has been explored (Pan et al., 2021; Wang et al., 2023). However, the substrate generation process of the N-end rule in intact cells, by modifying enzymes, such as deamidases and Arg transferase, is lacking.

**Figure 1.**
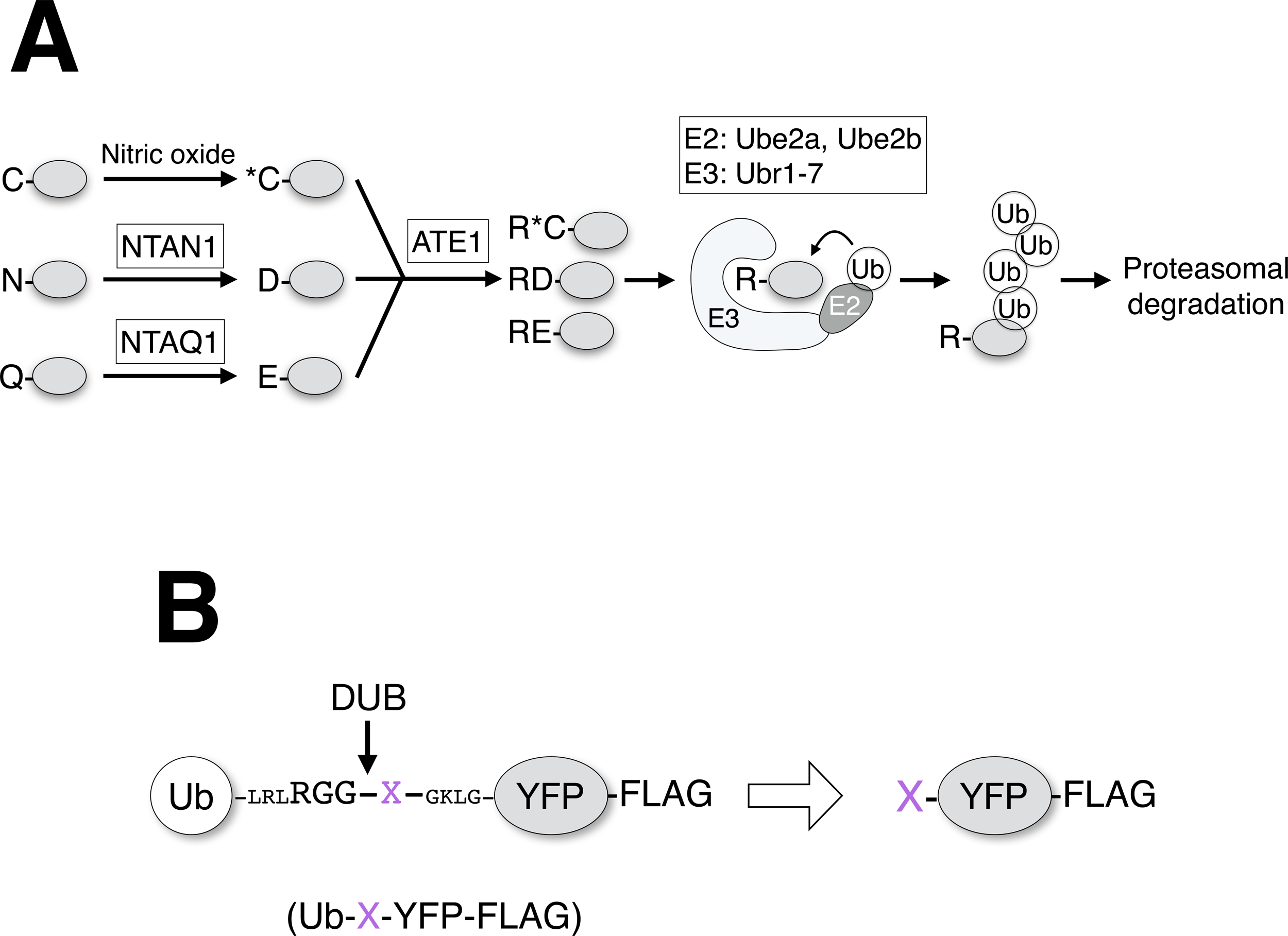
Schematic representation of the Arg-mediated N-end rule pathway and its artificial substrate, Ub-X-YFP-FLAG. (A) Proteins with Cys, Asn, or Gln at the N terminus are modified by NO, NTAN1, and NTAQ1, respectively, to oxidized cysteine (shown as an asterisk), Asp, and Glu. ATE1 adds an Arg residue to their N termini. E3 ligases (Ubr1-7) recognize these proteins having Arg at the N terminus as an N-end rule substrate. After the transfer of ubiquitin (Ub) by E2 (Ube2a or Ube2b), the substrate proteins are polyubiquitinated and targeted for proteasomal degradation. (B) Generation of the X-YFP-FLAG with the X amino acid at the N-terminus. Ub-X-YFP-FLAG contains a DUB-recognizing sequence motif (RGG) between the N-terminus Ub moiety and the C-terminus YFP-FLAG. After translation in cultured cells, cellular DUB processes Ub-X-YFP-FLAG at the RGG motif, and the resultant protein exposes the desired amino acid (shown as X in purple) at the N-terminus.

In this study, we explored the effects of N-terminal modification enzymes on the degradation of the N-end rule in cultured cell lines using artificial substrate proteins. We employed Ub-X-Yellow fluorescent protein (YFP)-FLAG, which is processed after ubiquitin (Ub), and the resultant protein contained an amino acid (described as X) at the N-terminus (Dantuma et al., 2000). We constructed M-, N-, D-, Q-, E-, and R-YFP-FLAG-expressing plasmids and verified that all these proteins, except for M-YFP-FLAG, were degraded by the N-end rule via the proteasome. N-terminal asparagine amidohydrolase 1 (NTAN1) and arginine tRNA protein transferase 1 (ATE) accelerate N-end rule degradation depending on their protein expression levels. Interestingly, N-terminal glutamine amidohydrolase 1 (NTAQ1) overexpression inhibited N-end rule degradation, which was enhanced by NTAQ1 siRNA. Additionally, the outer loop of NTAQ1 was found to be necessary for inhibiting Q-YFP-FLAG degradation, as predicted by the AlphaFold2 and MMseqs2 algorithms. Taken together, our results suggest that N-terminus-modifying enzymes of the N-end rule exhibit different effects on the degradation pathway in cultured cell lines. It is possible to regulate N-end rule degradation by the protein expression level of the deamidases by a mechanism that needs to be explored in more detail.

## Results

### Development of a cellular assay system for the N-end rule degradation

To investigate the details of Arg/N-end rule degradation within cells, we established an assay system with an exogenously expressed model substrate protein, Ub-X-YFP-FLAG. The Ub-X-YFP-FLAG undergoes processing by cellular deubiquitinases at the RGG motif between Ub and YFP moiety (Dantuma et al., 2000). Following excision, X-YFP-FLAG, which contains arbitrary amino acid at the N-terminus, is expressed (Figure 1B).

We focused our attention on the Asn and Gln routes of Arg/N-end rule degradation, as the Cys-mediated degradation route has been well characterized using endogenous substrates RGS4 and RGS5 (Heo et al., 2023; Hu et al., 2005; Lee et al., 2005). The protein expression of M-YFP-FLAG was not affected by proteasome inhibition (Figure 2A, lanes 1 and 2), as reported previously (Dantuma et al., 2000). In contrast, R-YFP-FLAG expression was significantly restored by a proteasome inhibitor, MG132 (Figure 2A, lanes 3 and 4). In transfected 293T cells, the fluorescence of R-YFP-FLAG increased when cells were treated with MG132 (Figure 2B, panels a and d), suggesting that YFP-FLAG bearing an N-terminal Arg is degraded by the proteasome.

**Figure 2.**
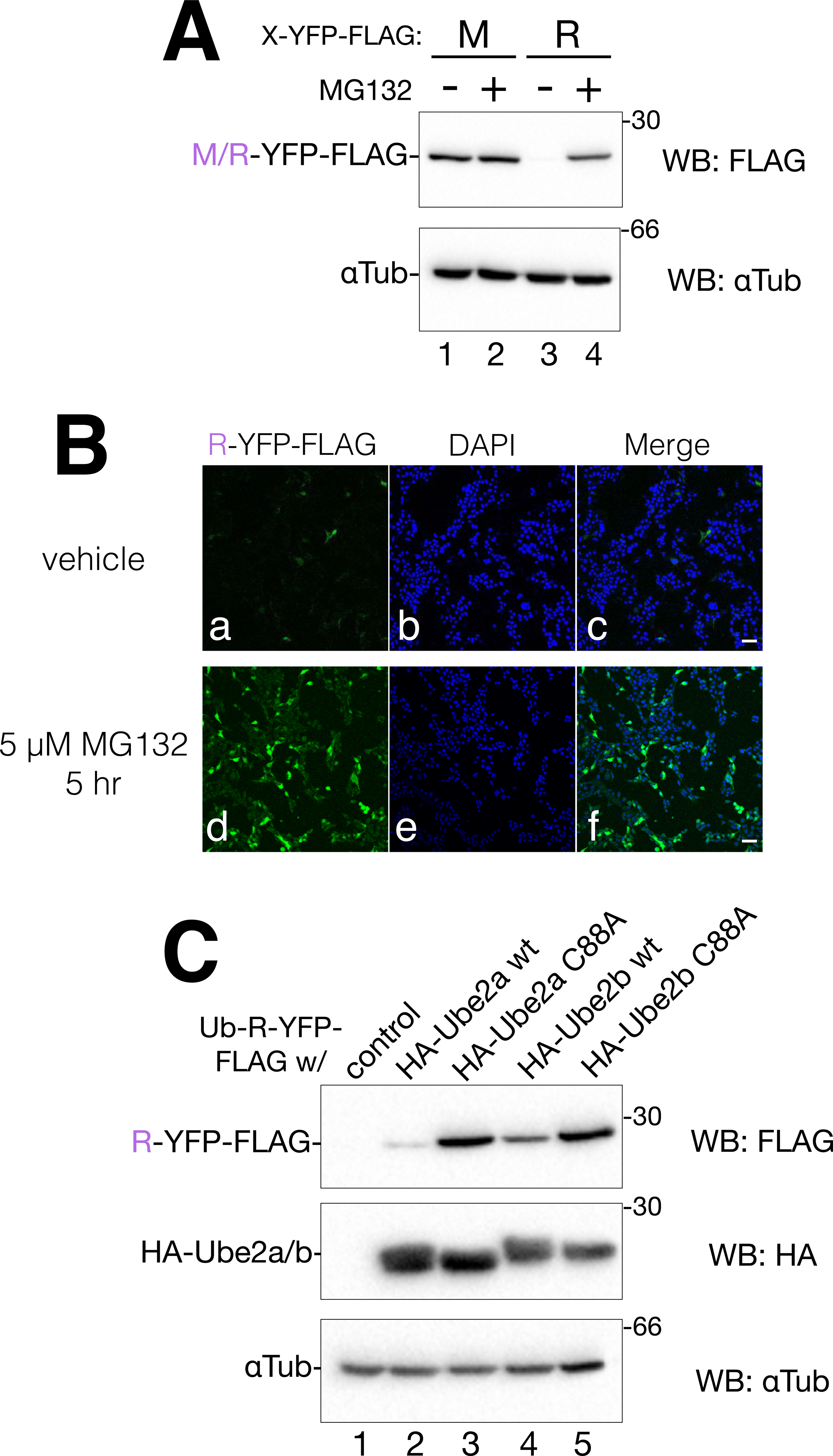
Arg-mediated N-end rule degradation in cultured cells. (A) 293T cells transfected with Ub-M-YFP-FLAG or Ub-R-YFP-FLAG were treated with or without 5 µM MG132 for 5 h. Western blotting was performed using the indicated antibodies. The target N-terminus amino acids are shown in purple. (B) Transfected 293T cells with Ub-R-YFP-FLAG grown on a coverslip were treated with vehicle (a-c) or 5 µM MG132 (d-f) for 5 h. After fixation and DAPI staining, confocal images were obtained. Merged images (scale bar 50 µm) are shown in panels c and f. (C) Ub-R-YFP-FLAG was co-transfected with the control or indicated vectors. Protein was detected after western blotting using the indicated antibodies.

Next, we constructed HA-tagged N-end rule E2s, Ube2a, and Ube2b, and co-expressed them with Ub-R-YFP-FLAG to examine the polyubiquitination before proteasomal degradation. We also verified the E2 activity of Ube2a and Ube2b using the point mutation C88A, which eliminates ubiquitin-binding activity and functions as a dominant-negative form (Siepmann et al., 2003). Both the C88A mutations of Ube2a and Ube2b significantly suppressed R-YFP-FLAG degradation (Figure 2C, lanes 3 and 5) compared to the co-expression of the control vector (Figure 2C, lane 1) or wild type (WT) (Figure 2C, lanes 2 and 4). Ube2a C88A mutation also inhibited R-YFP-FLAG degradation in HeLa cells (Supplemental Figure 1A). These results support the notion that our cellular assay system is useful for investigating the molecular mechanisms underlying N-end rule degradation.

### The expression level of NTAN1 affects N-YFP-FLAG degradation by the Arg/N-end rule pathway

We assessed whether the Arg/N-end rule pathway works in cultured cell lines and if the expression level of amidases affects the degradation. The deamidases NTAN1 and NTAQ1, responsible for converting Asn and Gln residues of the target proteins to Asp and Glu, respectively, have been identified and their structural analyses were performed (Park et al., 2014, Park et al., 2020). However, the actual cellular function of the Arg/N-end rule pathway in substrate generation is not well documented. Ub-N-YFP-FLAG, which exposes Asn at the N-terminus of YFP-FLAG, was constructed and co-expressed in 293T cells using an empty control plasmid, or NTAN1-HA. Overexpression of NTAN1-HA significantly enhanced the clearance of N-YFP-FLAG (Figure 3A, lanes 1-3). Upon co-expression of Ub-N-YFP-FLAG with HA-Ube2b C88A, a dominant-negative form of E2, the concentration of N-YFP-FLAG increased, indicating that N-YFP-FLAG is the target of Arg/N-end rule degradation (Figure 3A, lanes 1 and 4). This degradation of N-YFP-FLAG was also observed in HeLa cells (Supplemental Figure 1B).

**Figure 3.**
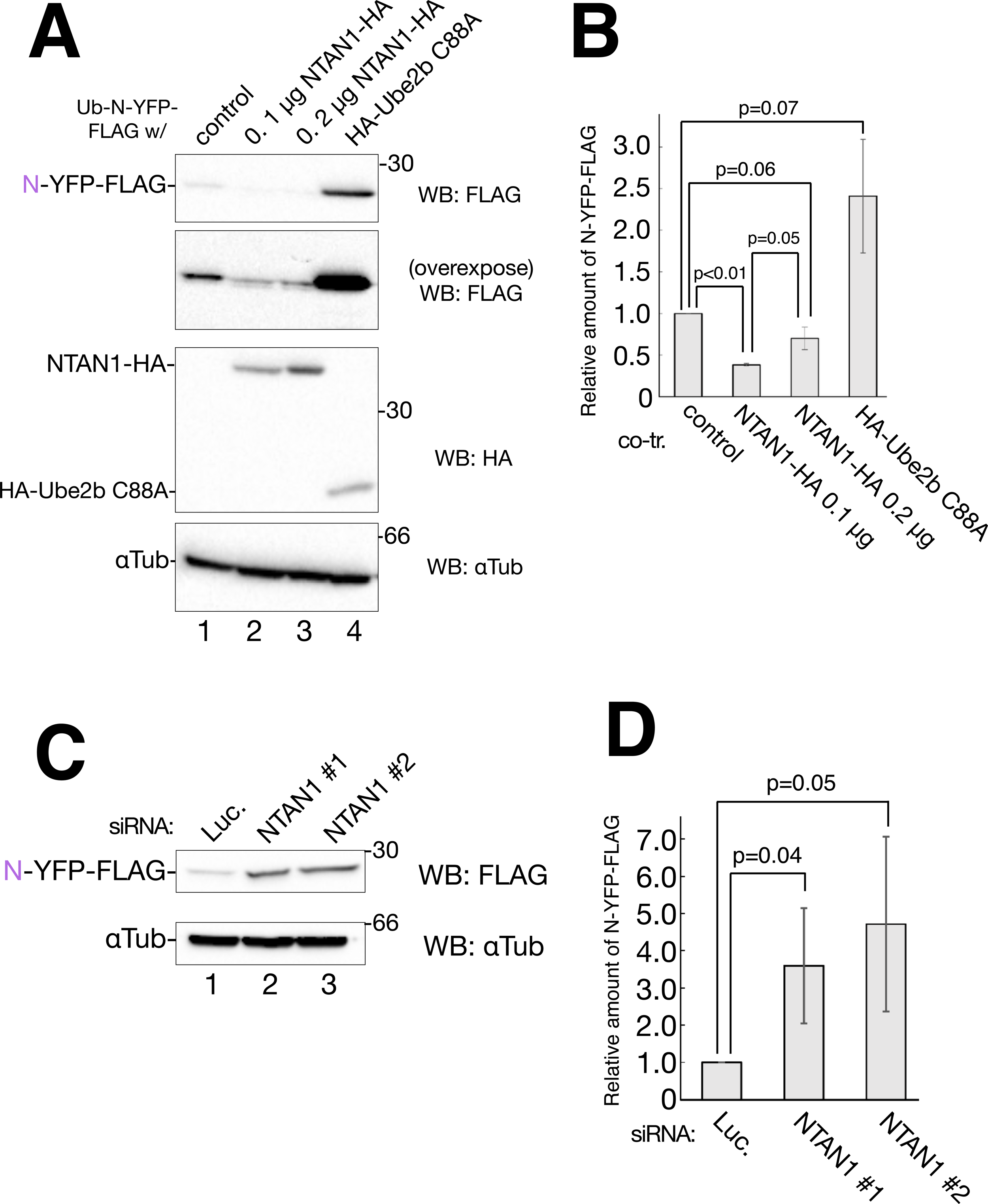
Effect of NTAN1 on Arg-mediated N-end rule degradation. (A) 293T cells were co-transfected with the Ub-N-YFP-FLAG plasmid (0.2 µg) and the control vector, NTAN1-HA, or HA-Ube2b C88A. After cell lysis and western blotting, proteins were detected using the indicated antibodies. (B) Quantification of A. The ratio was normalized with α-Tubulin, with samples of the control vector regarded to be 1.0. Error bars represent the standard deviation within the sample from the three independent experiments, corresponding *p*-values are indicated. (C) 293T cells were treated with siRNA against luciferase (Luc.) or NTAN1 and then transfected with Ub-N-YFP-FLAG. After cell lysis and western blotting, proteins were detected using the indicated antibodies. (D) Quantification of C. The ratio was normalized with α-Tubulin with samples of Luc considered to be 1.0. Error bars represent the standard deviation within the sample from the three independent experiments, corresponding *p*-values are indicated.

Next, we examined whether the protein expression level of NTAN1 is a primary factor in the degradation of N-YFP-FLAG. We downregulated the expression of NTAN1 mRNA using siRNA and confirmed its effect on the transfected NTAN1-HA (Supplemental Figure 2A), as the anti-NTAN1 antibody we obtained did not work in our experimental systems for endogenous NTAN1 (data not shown). As shown in Figure 3C and its densitometric analysis (Figure 3D), knockdown of the NTAN1 gene significantly increased the expression of N-YFP-FLAG. Taken together, these results suggest that NTAN1 promotes the deamidation of proteins containing N-terminal Asn residues for Arg/N-end rule degradation.

### ATE1 expression level promotes D-YFP-FLAG degradation via the Arg/N-end rule pathway

To obtain further insight into the Asn-mediated route of Arg/N-end rule degradation, we examined the effects of ATE1 on our assay system. Using the total RNA extracted from 293T cells, we successfully subcloned the ATE1 cDNA, which corresponds to the ATE1-2 isoform (referred to as ATE1), which can arginylate protein substrates containing Asp or Glu at the N-terminus (Saha et al., 2012). We confirmed the cytoplasmic localization of HA-tagged ATE1 using immunofluorescence experiments in HeLa cells (data not shown).

Co-expression of ATE1-HA with Ub-D-YFP-FLAG, which contains an N-terminal Asp residue, revealed that increasing ATE1 protein levels promoted D-YFP-FLAG degradation (Figure 4A, lanes 1 and 2). This enhancement was dependent on the amount of ATE1-HA protein expression (Figure 4B and C), and the expression of Ub-D-YFP-FLAG was restored by treatment with MG132 (Figure 4A, lanes 1 and 3). These results suggest that Arg transfer to the N-terminus of D-YFP-FLAG and subsequent proteasomal degradation occur in an ATE1 expression level-dependent manner.

**Figure 4.**
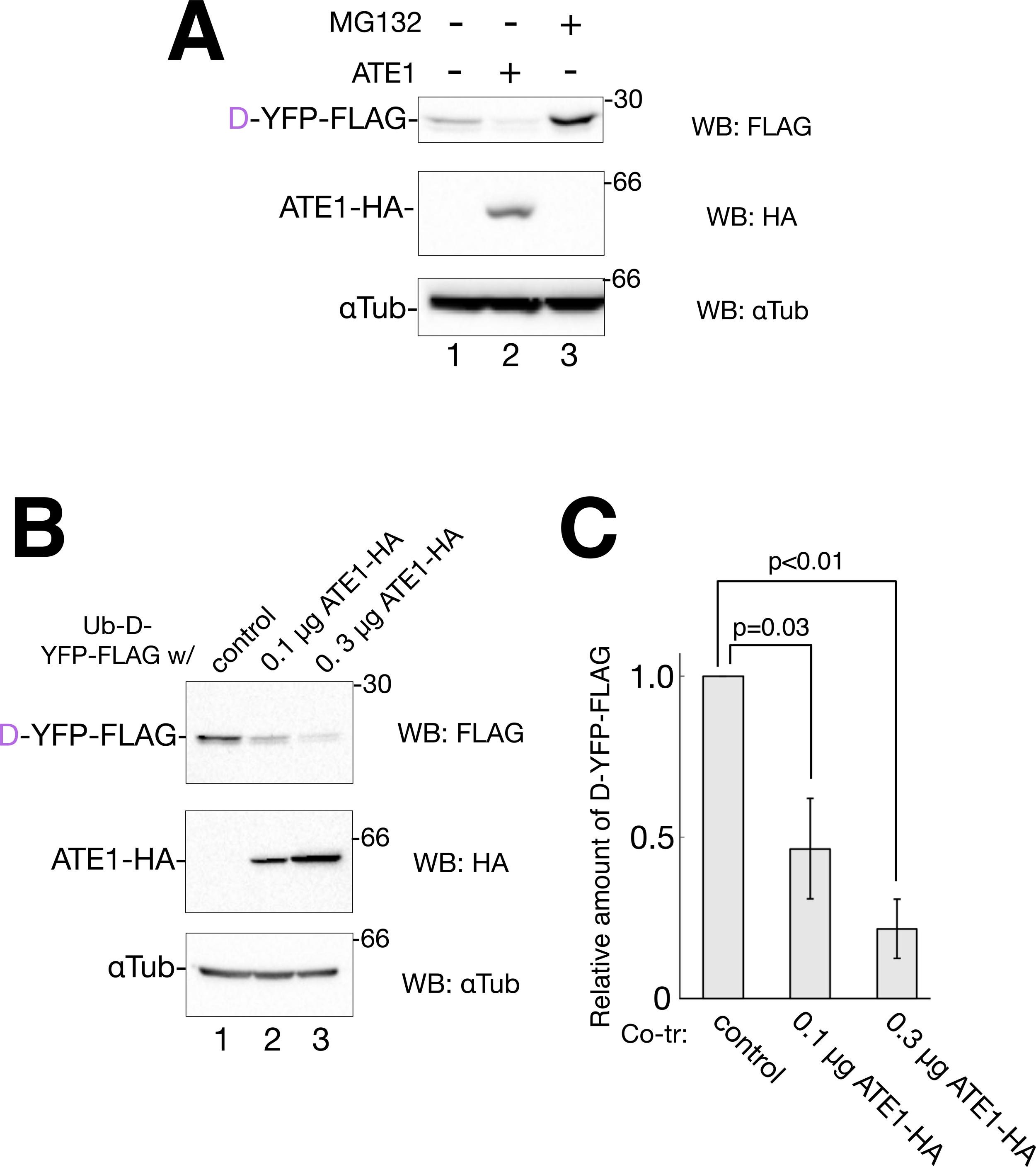
ATE1 promotes proteasomal degradation of D-YFP-FLAG. (A) 293T cells were transfected with Ub-D-YFP-FLAG plasmid (0.2 µg) and control vector (lane 1), 0.3 µg of ATE1 wt-HA (lane 2), or control vector with MG132 (lane 3). MG132 was added at a concentration of 1 µM after 32 h of transfection and incubated for 16 h. After cell lysis and western blotting, proteins were detected using the indicated antibodies. (B) Co-transfection of 293T cells with Ub-D-YFP-FLAG plasmid (0.2 µg) and control vector, 0.1 µg of ATE1 wt-HA, or 0.3 µg of ATE1 wt-HA was conducted for 48 hr. After cell lysis and western blotting, proteins were detected using the indicated antibodies. (C) Quantification of B. The ratio was normalized with α-Tubulin, with samples of the control vector regarded as 1.0. Error bars represent the standard deviation within the sample for the three independent experiments, corresponding *p*-values are indicated.

We further characterized ATE1 in HeLa cells using the ATE1 inhibitor, tannic acid, and point mutations (Supplemental Figure 3). Chemical screening revealed that tannic acid inhibited ATE1 (Kim et al., 2022; Saha et al., 2012). Co-transfection with Ub-D-YFP-FLAG and ATE1 and subsequent tannic acid treatment indicated that D-YFP-FLAG expression increased in a concentration-dependent manner (Supplemental Figure 3A). Analysis of the crystal structure of ATE1 obtained from the budding yeast *Kluyveromyces lactis* elucidated that ATE1 contains a zinc-finger motif comprising four cysteine residues (Kim et al., 2022). Hu et al. revealed that Cys71 and Cys72 of human ATE1 form an intramolecular disulfide bond induced by hemin binding (Hu et al., 2008). Furthermore, structural prediction using the AlphaFold2 algorithm suggested that the Cys71 residue of human ATE1 forms an intramolecular disulfide bond with Cys27 (data not shown). We introduced point mutations into the Cys71 residue of human ATE1 and examined the downstream Arg/N-end rule degradation. The replacement of Cys71 with Ala suppressed D-YFP-FLAG degradation (Supplemental Figure 3B, lanes 2 and 3). In contrast, ATE1 C71A was less expressed than WT ATE1 (Supplemental. 3B, lanes 2 and 3). These results suggest that in human ATE1, C71 contributes to its structural stability, potentially through zinc coordination or disulfide bonds. K419 of ATE1 is important for Arg transfer to the N-terminus of the Asp/Glu substrate protein, as demonstrated by the assays using *Saccharomyces cerevisiae* (Rai et al., 2006). Mutation of ATE1 K419 to Ala negatively affected D-YFP-FLAG degradation in our assay system (Supplemental Figure 3C). Collectively, ATE1 enhanced D-YFP-FLAG degradation via its Arg-transfer activity.

### Degradation of Q- and E-YFP-FLAG through the Arg/N-end rule pathway

Next, we used our cellular assay system to examine the Gln route of the Arg/N-end-rule pathway. Co-expression of ATE1 decreased the detection of Q-YFP-FLAG (Figure 5A, lanes 1 and 2). In addition, HA-Ube2b C88A and MG132 treatment increased Q-YFP-FLAG band intensity (Figure 5A, lanes 1, 3, and 4), suggesting that Q-YFP-FLAG serves as a target protein of the Arg/N-end rule degradation pathway, similar to N-YFP-FLAG. The expression level of E-YFP-FLAG was higher than that of Q-YFP-FLAG (Figure 5A, lanes 1 and 5), and Ube2b C88A and MG132 were less effective (Figure 5A, lanes 7 and 8 and Figure 5B, lanes 1 and 5), although ATE1 showed some effects on E-YFP-FLAG degradation (Figure 5A, lanes 5 and 6). However, there is no direct evidence explaining these results. The expression level of E-YFP-FLAG was not changed by the overexpression of NTAQ1 (Supplemental Figure 4, lanes 1 and 2), suggesting that NTAQ1 did not affect the turnover of its enzymatic product, E-YFP-FLAG. One interpretation for the lower degradation rate of E-YFP-FLAG involves sequential assistance through the association of ATE1 and NTAQ1, or transfer of the substrate proteins from NTAQ1 to ATE1, which could be required for efficient Arg transfer to E-YFP-FLAG by ATE1.

**Figure 5.**
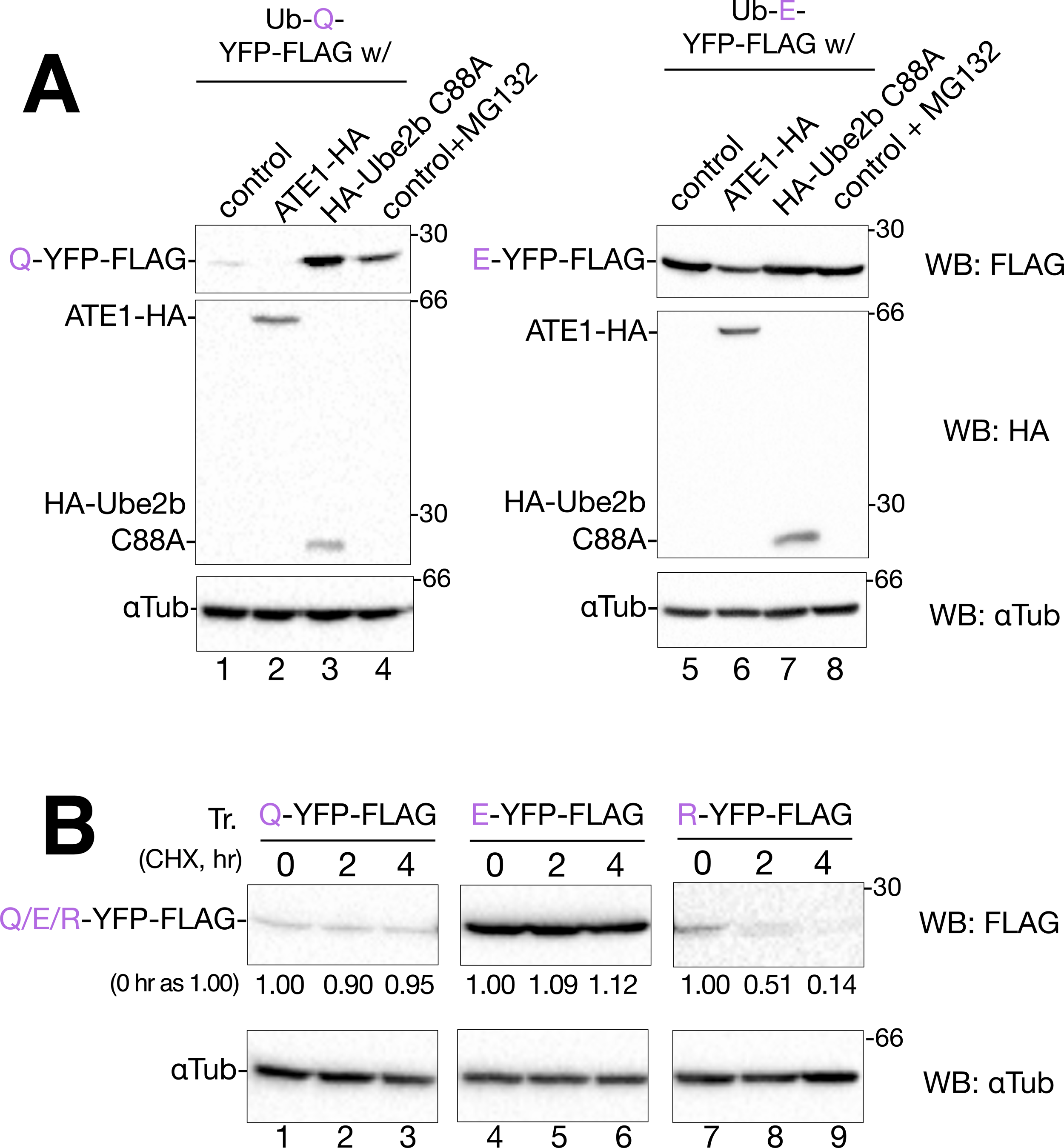
N-end rule degradation of Q, E, or R-YFP-FLAG. (A) 293T cells were transfected with 0.2 µg of Ub-Q-YFP-FLAG (lanes 1-4) or Ub-E-YFP-FLAG (lanes 5-8) plasmid for 48 h. Co-transfection with the control vector (lanes 1, 4, and 5, 8), ATE1 wt-HA (lanes 2 and 6), or HA-Ube2b C88A (lanes 3 and 7) was conducted. MG132 was added at a concentration of 1 µM after 32 h of transfection and incubated for 16 h (lanes 4 and 8). Proteins of interest were detected by immunoblotting using the indicated antibodies. (B) 293T cells were transfected with 0.2 µg of Ub-Q-YFP-FLAG (lanes 1-3), Ub-E-YFP-FLAG (lanes 4-6), or Ub-R-YFP-FLAG (lanes 7-9). After 48 h, transfected cells were incubated with fresh media containing 10 µg/mL CHX for the indicated time. Proteins of interest were detected using immunoblotting with the indicated antibodies. Values of Q, E, and R-YFP-FLAG are normalized to those of α-Tubulin and the time of each zero point was set at 1.00.

To further characterize the Gln-mediated Arg/N-end rule pathway, a cycloheximide (CHX)-chase assay was conducted in transfected cells. Although the turnover rates of Q-YFP-FLAG and E-YFP-FLAG were slow (Figure 5B, lanes 1–6), the degradation rate of R-YFP-FLAG was faster (Figure 5B, lanes 7–9). These results suggest that the conversion of the N-terminal Gln to Arg via Glu is the rate-limiting step in the Arg/N-end rule.

### The expression level of NTAQ1 negatively affects Q-YFP-FLAG degradation by the Arg/N-end rule pathway

To verify that the deamidation and subsequent degradation of Q-YFP-FLAG were dependent on the protein expression of NTAQ1, Ub-Q-YFP-FLAG was co-expressed with NTAQ1-HA in 293T cells. Notably, the amount of Q-YFP-FLAG increased depending on the expression level of NTAQ1-HA (Figure 6A). This suggests that overexpression of NTAQ1-HA suppresses the degradation of Q-YFP-FLAG, in contrast to that observed for N-YFP-FLAG and NTAN1-HA (Figure 3A). The degradation rate of Q-YFP-FLAG co-transfected with the control vector or NTAQ1-HA did not show a significant change in CHX-chase experiments (Figure 6B), which was consistent with the results shown in Figure 6A.

**Figure 6.**
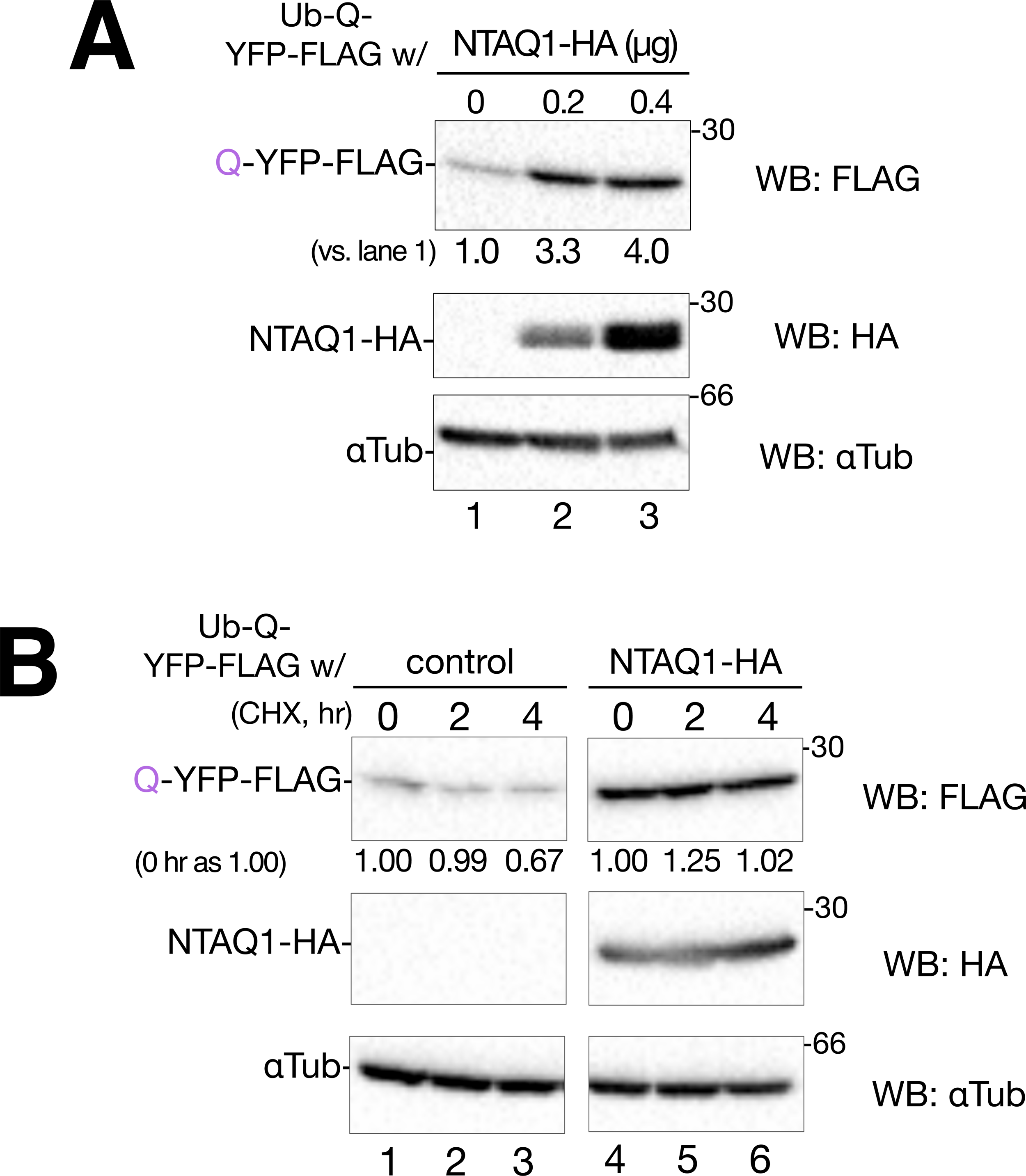
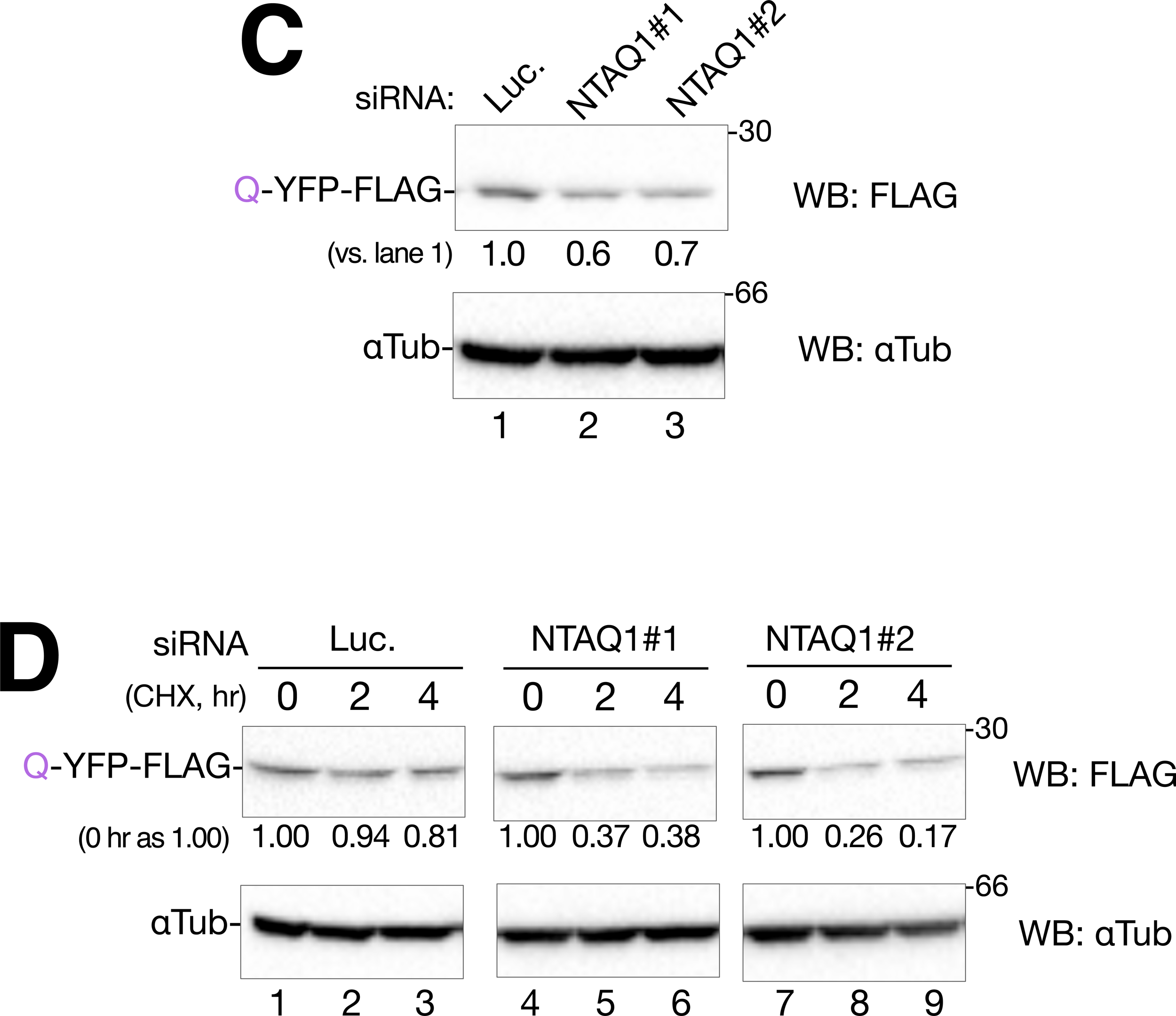
Expression of NTAQ1 negatively affects Q-YFP-FLAG degradation. (A) Co-transfection of 293T cells with the Ub-Q-YFP-FLAG plasmid (0.2 µg) and control vector, 0.2 or 0.4 µg of NTAQ1 wt-HA was conducted for 48 h. After cell lysis and western blotting, proteins were detected using the indicated antibodies. Values of Q-YFP-FLAG are normalized to those of α-Tubulin and the co-transfection of the control vector (lane 1) was set at 1.0. (B) 293T cells were co-transfected with 0.2 µg of Ub-Q-YFP-FLAG and the control vector (lanes 1-3) or NTAQ1 wt-HA (lanes 4-6), respectively. After 48 hr, transfected cells were incubated with fresh media containing 10 µg/mL CHX for the indicated time. Proteins of interest were detected using immunoblotting with the indicated antibodies. Values of Q-YFP-FLAG are normalized to α-Tubulin and the time of each zero point was set as 1.00. (C) 293T cells were treated with siRNA against luciferase or NTAQ1 and then transfected with Ub-Q-YFP-FLAG. After cell lysis and western blotting, proteins were detected using the indicated antibodies. Values of Q-YFP-FLAG are normalized to those of α-Tubulin and the control siRNA was set at 1.0. (D) SiRNA and subsequent transfection of 293T cells were conducted as mentioned in C. The cells were incubated with fresh media containing 10 µg/mL CHX for the indicated time. Proteins of interest were detected using immunoblotting with the indicated antibodies. Values of Q-YFP-FLAG are normalized to those of α-Tubulin and the time of each zero point was set at 1.00.

We downregulated the expression of the NTAQ1 gene via siRNA targeting and subsequently reduced protein expression to examine the Q-YFP-FLAG degradation. The anti-NTAQ1 antibody we purchased did not work in our experimental system (data not shown). Therefore, the siRNA effect was verified by analyzing NTAQ1-HA expression levels using western blotting with an anti-HA antibody, which confirmed the significant downregulation of NTAQ1-HA (Supplemental Figure 2B). In NTAQ1-knockdown 293T cells, the remaining Q-YFP-FLAG was decreased compared to that in the negative siRNA control (Figure 6C). Furthermore, NTAQ1 knockdown accelerated the rate of Q-YFP-FLAG degradation (Figure 6D). These results suggest that high levels of NTAQ1 negatively affect Q-YFP-FLAG degradation within cells.

### NTAQ1 overexpression reduces polyubiquitination of Q-YFP-FLAG

In the Gln route of the Arg/N-end rule pathway, three protein modification steps are required: deamidation of the N-terminus of Gln by NTAQ1, transfer of Arg to the N-terminus by ATE1, and E2-E3 mediated polyubiquitination (Varshavsky, 2019). To understand the negative effects of NTAQ1 overexpression, the polyubiquitination of Q-YFP-FLAG was examined.

Ub-Q-YFP-FLAG was increased by the overexpression of NTAQ1-HA (Figure 7, lanes 2 and 3 in the total cell lysate) or MG132 treatment (Figure 7, lanes 2 and 4 in the total cell lysate), as demonstrated in Figures. 5 and 6. Whole cellular ubiquitinated proteins, detected by an antibody recognizing K48-polyubiquitinated proteins destined for proteasomal degradation, showed an increase only by MG132 treatment (Figure 7, lane 4), suggesting that the expression of Ub-Q-YFP-FLAG or NTAQ1-HA did not affect overall cellular polyubiquitination. By isolating FLAG-tagged proteins with anti-FLAG agarose beads, Q-YFP-FLAG was found to be polyubiquitinated (Figure 7, lane 6), and MG132 treatment increased the amount of polyubiquitinated Q-YFP-FLAG (Figure 7, lane 8), suggesting that polyubiquitinated Q-YFP-FLAG accumulated owing to proteasomal inhibition. In the co-expression of NTAQ1-HA, polyubiquitinated Q-YFP-FLAG was decreased compared to that in the control vector co-transfection (Figure 7, lanes 6 and 7). These results indicate that the overexpression of NTAQ1-HA did not affect the proteasome activity. Rather, the overexpressed NTAQ1-HA negatively affects the deamidation by NTAQ1, the arginylation by ATE1, or the polyubiquitination by the E2-E3 complex.

**Figure 7.**
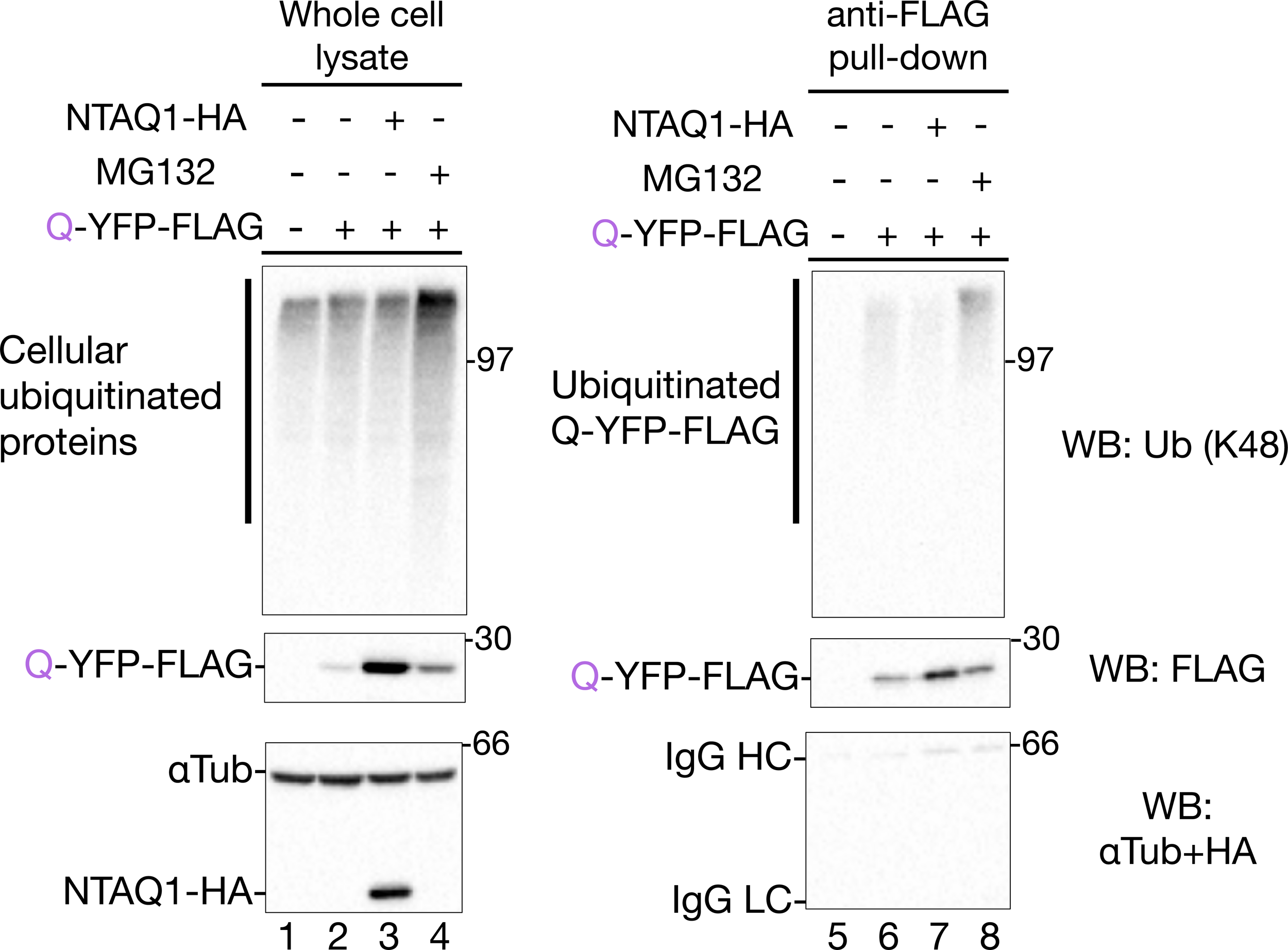
Polyubiquitination of Q-YFP-FLAG is reduced by NTAQ1 293T cells were transfected with the control vector (lanes 1 and 5), Ub-Q-YFP-FLAG (lanes 2, 4, 6, and 8), or Ub-Q-YFP-FLAG and NTAQ1 wt-HA (lanes 3 and 7). In lanes 4 and 8 samples, MG132 was added at a concentration of 1 µM at 32 h after transfection and incubated for 16 h. Cell lysates were separated for use as whole-cell lysates (lanes 1–4) or anti-FLAG pull-downs (lanes 5–8). After extraction of FLAG-tagged proteins using the FLAG peptide and reducing SDS-PAGE, the proteins of interest were detected using the indicated antibodies. For western blot analysis of αTub+HA, Ub, and FLAG 1.7%, 3.3%, and 5% of total cell lysate were used as input, respectively. For anti-FLAG pull-down, 22.5%, 22.5%, and 45% of total cell lysate were used for the western blot analysis of a mix of αTub and HA (αTub+HA), Ub, and FLAG.

### The outer loop of NTAQ1 is critical for Q-YFP-FLAG accumulation

It hypothesized that if NTAQ1 negatively affects ATE1 activity, these proteins should be physically associated with the Arg/N-end rule degradation. To inspect the functional interaction between ATE1 and NTAQ1, we conducted immunoprecipitation experiments in which pull-down of FLAG-tagged ATE1 was initially performed, followed by immunoblotting using anti-HA for NTAQ1-HA. However, this interaction was not observed under our experimental conditions (data not shown). Oh et al. revealed that the Arg/N-end rule pathway factors form a functional complex comprising E3 Ubr1, E2 Ube2a/b, NTAN1, NTAQ1, and ATE1, using various binding assays (Oh et al., 2020). They reported an association of NTAQ1 with Ubr1 and NTAN1, but not with ATE1 (Oh et al., 2020).

We predicted the NTAQ1-ATE1 complex formation using the machine-learning-driven protein-structure prediction software AlphaFold2.0 (Jumper et al., 2021), and ColabFold v1.5.5: AlphaFold2 using MMseqs2 for complex analysis (Mirdita et al., 2022). The prediction of complex formation by MMseqs2 suggested two association points between ATE1 and NTAQ1 (Figure 8A, Contact sites 1 and 2). A detailed analysis of the interatomic contacts was performed using the Residue Interaction Network Generator (RING) server, version 3.0 (Clementel et al., 2022). As predicted by the RING analysis, S152, S153, and R157 in the outer loop of NTAQ1 (contact site 1) appeared to approach ATE1 (Supplemental Figure 5A and B). The crystal structure analysis of NTAQ1 revealed that residues 150–170 of NTAQ1 form an outer loop (Park et al., 2014). The X-ray structures of human NTAQ1 (Park et al., 2014) and NTAQ1 in complex with ATE1 by AlphaFold2 and MMseqs2 modeling showed good overall structural agreement (Supplemental Figure 5C), suggesting that the structure of NTAQ1 did not change drastically when associated with ATE1. We constructed a 151–166 deletion mutant of NTAQ1 and co-expressed it with Ub-Q-YFP-FLAG. Compared to the control vector and WT NTAQ1-HA (Figure 8B, lanes 1 and 2), this outer loop-deleted NTAQ1 (NTAQ1 Δ151-166) did not exhibit an inhibitory effect against Q-YFP-FLAG (Figure 8B, lane 3). In addition, this deletion did not affect E-YFP-FLAG degradation (Supplemental Figure 4). These results suggest that NTAQ1 Δ151-166 affects only the deamidation of Q-YFP-FLAG and not the subsequent arginylation of ATE1 or WT NTAQ1. We also attempted to identify other point mutations in NTAQ1, including R59 and R70, in the contact site 2. However, these NTAQ1 mutants did not recover from Q-YFP-FLAG degradation (data not shown), suggesting that these residues are not involved in the inhibitory effect of NTAQ1. These results indicate that residues 151–166 of NTAQ1, the region of contact site 1, are important for inhibiting Q-YFP-FLAG degradation through the Arg/N-end rule pathway. Collectively, our results using the cellular assay systems revealed that the Asn- and Gln-mediated Arg/N-end rule pathway is regulated by NTAN1, NTAQ1, and ATE1, which are dependent on the expression level of the deamidases in different regulatory mechanisms.

**Figure 8.**
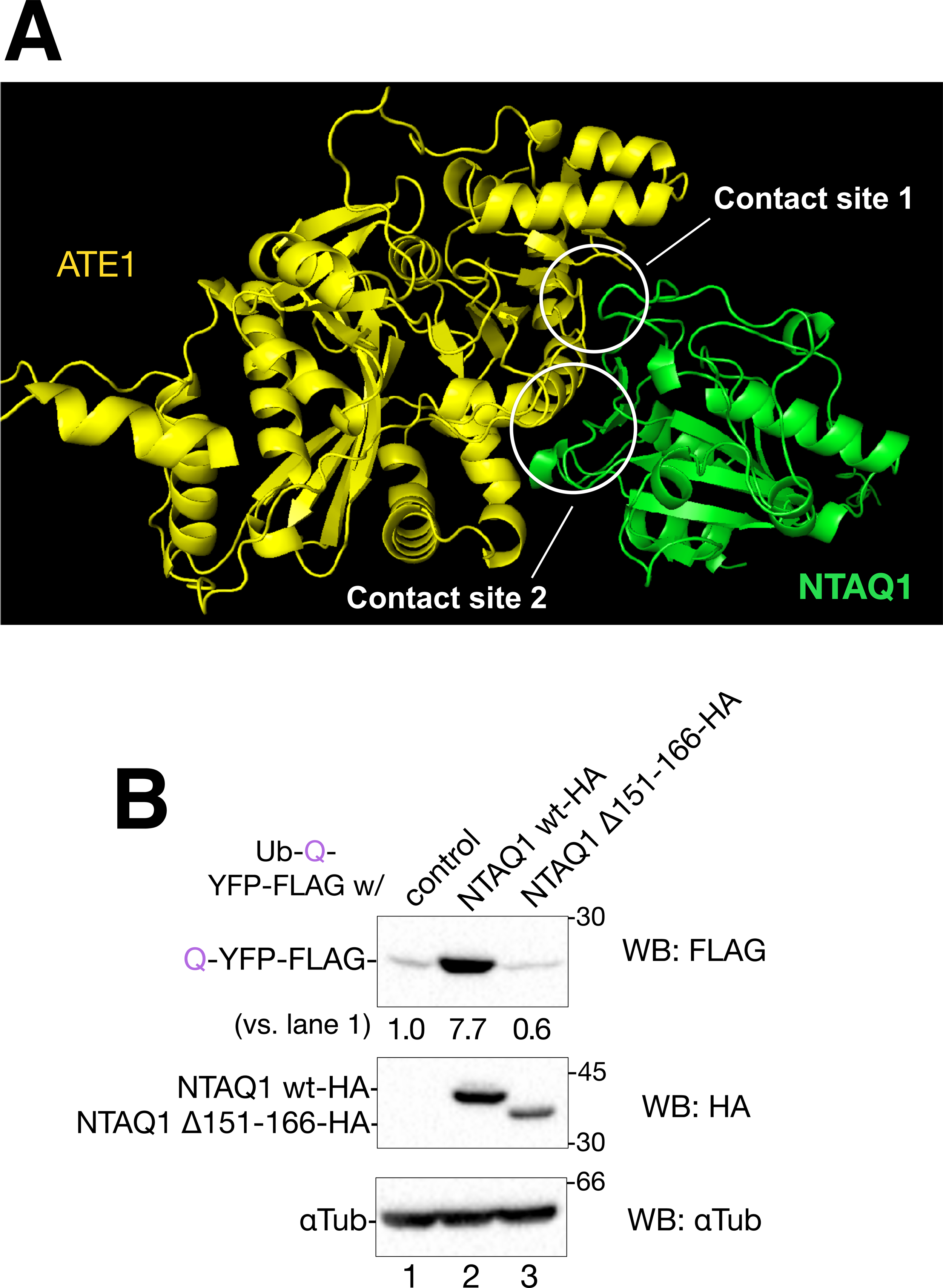
Recovery of Q-YFP-FLAG degradation by deletion of the NTAQ1 151–166 outer loop region. (A) Complex formation and interaction sites of human NTAQ1 (green) and ATE1 (yellow) were predicted using AlphaFold2 and MMseqs2. The inferred region of the interaction of NTAQ1 and ATE1 is shown as contact sites 1 and 2. (B) 293T cells were co-transfected with 0.2 µg of Ub-Q-YFP-FLAG and the control vector (lanes 1), NTAQ1 wt-HA (lane 2), or NTAQ1 Δ151–166 HA (lane 3), respectively. After 48 h transfection, cell lysates were prepared, and SDS-PAGE was conducted. Proteins of interest were detected using immunoblotting with the indicated antibodies. Values of Q-YFP-FLAG are normalized by those of α-Tubulin and the control sample was set at 1.0.

## Discussion

The N-end rule degradation occurs from single cellular organisms to vertebrates and the N-end rule factors are expressed in almost all human tissues. Additionally, in bacteria, the ClpS-ClpAP protease complex targets proteins containing non-Met at their N-terminus (Erbse et al., 2006; Tobias et al., 1991). Therefore, modification of the N-terminus of proteins to non-Met amino acids is considered a universal signal for protein degradation. The molecular mechanism of the N-end rule has been extensively studied and well-characterized using either a yeast assay system or an in vitro polyubiquitination assay (Erbse et al., 2006; Tobias et al., 1991). In the present study, we established a cellular assay system using an artificial substrate construct, Ub-X-YFP-FLAG, and verified the role of two amidases, NTAN1 and NTAQ1, and an Arg-transferase, ATE1, in the N-end rule degradation mechanism.

Several studies have reported that NTAN1 and NTAQ1 play an important role in eukaryotic activity. NTAN1 is a well-characterized N-end rule gene that is conserved across yeast and humans (Baker & Varshavsky, 1995; Grigoryeve et al., 1996). NTAN1 knockout mice exhibit altered activity, social behavior, and spatial memory (Kwon et al., 2000), and NTAN1 expression was observed to be downregulated upon the conversion of myoblasts into myotubes (Grigoryeve et al., 1996). Furthermore, Dubnau et al. reported that a mutation in the *Drosophila NTAQ1* gene causes defects in long-term memory (Dubnau et al., 2003). Genetic defects in both NTAN1 and NTAQ1 cause severe brain function phenotypes, suggesting a role for the Arg/N-end rule pathway in brain development and its functional regulation.

Understanding the importance of N-end rule degradation in physiological processes has increased, although the types of proteins targeted by the N-end rule remain an enigma. MAP2 and human tropomyosin-receptor kinase-fused gene (TFG) are endogenous NTAN1 substrates (Hirai et al., 2006; Nguyen et al., 2019). We assessed the TFG degradation through the Arg/N-end rule pathway using NTAN1-HA and FLAG-tagged TFG in our assay system. However, the effect of NTAN1-HA overexpression on TFG protein expression was not significant (data not shown). We assume that this relies on the method used to expose the Asn residue at the N-terminus. In the Ub-X-YFP-FLAG expression framework, X-YFP-FLAG, which contains X amino acids at the N-terminus, generates only proteins with X amino acids at the N-terminus. The first Met in TFG can be acetylated by the N-terminal acetyltransferase complex NatB since it contains Asn at the second amino acid position (Drazic et al., 2016). In more than half of newly synthesized proteins, Met is removed by methionine aminopeptidases (Bradshaw et al., 1998). However, because the activity of the aminopeptidase depends on the sequence, not all of the first Mets are successfully removed from the newly synthesized proteins by aminopeptidases (Xiao et al., 2010). Therefore, it is necessary to understand the mechanism by which substrates are generated and presented by the Arg/N-end rule in cells when the target protein contains an N-degron at the second amino acid.

The NTAQ1 gene is not present in all eukaryotic cells except the budding yeast *Saccharomyces cerevisiae* (Wang et al., 2009). In yeast, NTAN1p displays a polyvalent function that deamidates not only the protein containing the N-terminus Asn but also Gln (Baker & Varshavsky, 1995). Therefore, it is hypothesized that the types of N-end rule substrate proteins have diversified during evolution and that NTAQ1 might be added to the deamidase repertoire to accommodate the spectrum of N-end rule substrates. Similar cases have also been elucidated for ER protein quality control systems. A type I transmembrane protein calnexin is a lecint-like chaperone and predominantly interacts with transmembrane client proteins. Calnexin is found in both *Saccharomyces cerevisiae* and *Schizosaccharomyces pombe,* while its soluble ortholog, calreticulin, is only found in *Saccharomyces cerevisiae*. Based on their topological differences and biochemical evidence, an assigned task was proposed: calnexin preferentially associates with N-glycans in membrane-closed regions, while calreticulin associates with N-glycans away from the membrane (Hebert et al., 1997), or may serve as a backup for calnexin (Wada et al., 1995). Moreover, EDEM1, which promotes the degradation of misfolded ER proteins (ER-associated degradation, ERAD), exists in both soluble and transmembrane forms. In ERAD processes, soluble EDEM1 prefers soluble ERAD substrates and ERAD-related factors, while transmembrane-form EDEM1 prefers membrane-anchored ERAD substrates and ERAD factors (Tamura et al., 2011). These might serve as good examples of how the protein quality control mechanisms have evolved.

Because the prediction of the ATE1-NTAQ1 interaction suggested that three hydrogen bonds were concentrated around the outer loops 151–166, it is possible that NTAQ1 associates with ATE1 via this region (Supplemental Figure 5A and B). The deletion of the outer loop of NTAQ1 abrogated the negative effects of NTAQ1 overexpression on Q-YFP-FLAG degradation (Figure 8B). Although these results suggest a functional association between ATE1 and NTAQ1 during the Arg/N-end rule substrate generation, it is necessary to demonstrate the intact interaction of ATE1 and NTAQ1 using highly sensitive techniques such as TurboID (Branon et al., 2018) or the fluorescence-based proximity ligation assay (Hegazy et al., 2020). In this study, we revealed that the expression levels of NTAN1 and NTAQ1 positively and negatively affect the degradation of artificial substrates in concert with ATE1, respectively, via the Arg/N-end rule. Our results propose a negative feedback-like regulation of NTAQ1 in which the protein amount of NTAQ1 is strictly regulated in cells to prevent the overactivation of the Gln route of the Arg/N-end rule degradation by a transcriptional or protein level. In future studies, comprehensive identification of endogenous Arg/N-end rule substrates and elucidation of the physiological role of Arg/N-end rule degradation are required.

## Materials and Methods

### Plasmids

A plasmid expressing Ub-X-YFP-FLAG was constructed as follows: (1) the pHA-Ub-YFP: YFP moiety of pEYFP-N1 (Clontech, Mountain View, CA, USA) was amplified using PCR with primers HA-Ub-R-YFP f and r. HA-Ub-R-GFP was obtained from Integrated DNA Technologies (Coralville, IA, USA), and the YFP cDNA was assembled using an In-fusion HD cloning kit (Takara-Bio, Shiga, Japan). (2) pCX4-bsr-Ub-R-YFP: The Ub-R-YFP fragment was obtained from pHA-Ub-YFP via PCR amplification with primers Ub-R-YFP in-fusion f and r. Simultaneously, vector pCX4-bar was linearized through PCR amplification using primers pCX4bsr in-fusion f and r. These two fragments were assembled using In-fusion HD. (3) pCX4-bsr-Ub-R-YFP-FLAG: The FLAG fragment was obtained via PCR amplification of pCX4-bsr-EDEM1-FLAG (Miura et al., 2023) using the primers EDEM1-FLAG C term f and pCX4bsr 3006r. pCX4-bsr-Ub-R-YFP was linearized via PCR using the primers pCX4bsr3006f and Ub-R-YFP-FLAG r. The fragments were assembled using NEBuilder (New England Biolabs, Ipswich, MA, USA).

Plasmids expressing HA-Ube2a WT, HA-Ube2a WT, NTAN1-HA, NTAQ1-HA, and ATE1-HA constructed for this study were generated through reverse transcription using total RNA extracted from 293T, followed by PCR, and cloned into pCMV-HA-N or pCMV-HA-C (Clontech). Notably, we obtained an ATE1 DNA clone of the ATE1-2 isoform (Uniprot ID: O95260-2) from the total RNA of 293T cells. Point mutations at C88A in HA-Ube2a, C88A in HA-Ube2b, and C71A and K419A in ATE1-HA were introduced using standard inverse PCR with the corresponding mutagenic primers. The Ub-M/N/D/Q/E/-YFP-FLAG expression vector from pCX4-bsr-Ub-R-YFP-FLAG was constructed by standard inverse PCR with the corresponding mutagenic primers. Deletion of amino acids 151–166 of the NTAQ1-HA expressing plasmid was conducted through reverse PCR. Plasmid DNA sequences were confirmed using Sanger sequencing. All primers used in this study are listed in Supplemental Table 1.

### Antibodies and chemicals

Triton X-100, E64 (an antipain agent), tannic acid, and N-(2-hydroxy ethyl)-piperazine ethanesulfonic acid (HEPES) were purchased from Nacalai Tesque (Kyoto, Japan). 4!,6-diamidino-2-phenylindole (DAPI) was purchased from FUJIFILM Wako, Japan. MG132 was obtained from Peptide Institute Inc. (Osaka, Japan). Leupeptin, pepstatin, Mowiol, CHX, FLAG peptide (Sigma, F4799), and anti-FLAG agarose beads (Sigma, A2220) were purchased from Sigma Aldrich (St. Louis, MO, USA). Mouse anti-FLAG (Sigma, F1804), mouse anti-HA (MBL International, M180-3), mouse anti-α-Tubulin (Cedarlane, CLT9002), and rabbit monoclonal anti-UbK48 (Millipore, 51307) antibodies were used at a 1:4000, 1:4000, 1:10000 or 1:1000 dilution for western blotting.

### Cell culture, transfection, siRNA, and drug treatment

HEK293T and HeLa cells (kindly provided by Dr. Ikuo Wada, Fukushima Medical University, Fukushima, Japan) were cultured at 37 °C in 5% CO_2_ with humidity. The cells were maintained in Dulbecco!s modified Eagle!s medium (DMEM; Invitrogen, Carlsbad, CA, USA) containing 10% fetal bovine serum (FBS; Invitrogen), 1 mM L-glutamine (Nacalai Tesque), and 1% penicillin/streptomycin (Nacalai Tesque). The cells were authenticated using a Universal Mycoplasma Detection Kit (e-MycoTM, iNtRON, South Korea).

Plasmid transfection of 293T and HeLa cells was performed using polyethyleneimine (Polysciences Inc., Warrington, PA, USA) and HilyMax (DOJINDO Laboratories, Kumamoto, Japan), respectively, according to the manufacturer!s instructions. Ub-X-YFP-FLAG expressing vectors (0.2 µg) were used with the pcDNA 3.1 vector (Invitrogen) as the control vector to adjust the total plasmid amount by 0.5 µg per 35 mm dish. Gene knockdown by siRNA was performed using RNAiMAX (Invitrogen) according to the manufacturer!s protocol. Cells were incubated with 10 nM siRNA (Japan Bio Services Co., Ltd., Saitama, Japan) for 72 h.

Cell lysis and subsequent biochemical experiments were performed as previously described (Miura et al., 2023). Before lysis, the cells were washed once with PBS and incubated on ice with PBS containing 10 mM *N*-ethyleneimine (Kanto Chemical Co., Inc., Tokyo, Japan) for 10 min to modify the free thiol groups of Cys and prevent further modification. Subsequently, cells were solubilized in the lysis buffer containing 1% Triton X-100, 20 mM HEPES, pH=7.5, 150 mM NaCl, and protease inhibitor mix (5 µg/mL E64, antipain, leupeptin, and pepstatin). The lysate was centrifuged at 4 °C, 18,000 × *g* for 10 min to obtain a soluble supernatant fraction. Following the addition of sodium dodecyl sulfate-polyacrylamide gel electrophoresis (SDS-PAGE) sample buffer containing 0.1 M dithiothreitol, the proteins were denatured at 90 °C for 5 min. To analyze protein turnover, transfected 293T cells were treated with culture media containing 10 µg/mL CHX for a time period as indicated in each figure legend.

### Western blotting and immunoisolation

After SDS-PAGE, proteins were transferred onto polyvinylidene difluoride (PVDF) membranes (Immobilon; Millipore, Burlington, MA, USA). Incubation with primary antibodies was carried out at 25 °C for 1h. Treatment of horseradish peroxidase-conjugated goat anti-rabbit or mouse secondary antibodies (Sigma-Aldrich) was conducted at 25 °C for 30 min. Protein bands of interest were detected using a chemiluminescent reagent (luminol, *p*-coumaric acid, and hydrogen peroxide) and a ChemiDoc system (Bio-Rad, Hercules, CA, USA). Densitometric analysis was performed using ImageJ software v1.53 (National Institute of Health, Bethesda, MD, USA).

The expression of proteins of interest in the western blotting experiment was normalized to that of α-Tubulin as a cellular loading control. In all western blotting images, the proteins of interest and the molecular weights of the marker proteins are indicated on the left and right sides of the blot, respectively. The lane numbers are shown at the bottom. Data are shown as the mean of at least three independent experiments and were compared with the control using a two-tailed Student!s t-test. Error bars represent the standard deviation between experiments.

FLAG-tagged proteins were immunoisolated as previously described (Miura et al., 2023). FLAG-tagged proteins isolated with anti-FLAG M2 agarose beads were eluted following incubation with 50 mM Tris-HCl (pH 7.5) containing 150 mM NaCl and 500 µM FLAG peptide at 25 °C for 15 min with gentle rotation.

### Confocal microscopy

293T cells transfected with Ub-R-YFP-FLAG were grown on coverslips in a 35 mm dish for 48 h and then incubated with or without 5 µM MG132 for 5 h. Cells were washed twice with PBS and fixed with 4% paraformaldehyde in PBS (FUJIFILM Wako) at 25 °C for 10 min. After staining with 1/5000 DAPI at 25 °C for 3 min, the coverslips were rinsed with distilled water and mounted face-down on glass slides using Mowiol. Fluorescent YFP and DAPI stained images were obtained using a confocal laser microscope with an objective lens at a magnification of 20× (LSM 780, Carl Zeiss, Inc., Oberkochen, Germany).

### Complex structure prediction

The structure of the human ATE1 and NTAQ1 complex was predicted using ColabFold v1.5.5 MMseqs2 with AlphaFold2 (Mirdita et al., 2022). The interchain atomic interactions of the predicted ATE1-NTAQ1 complex were visualized and evaluated using RING software version 3.0 (Clementel et al., 2022). A comparison of the NTAQ1 structure between the crystal structure (PDB: 4w89) and the AlphaFold2-generated structure was performed and visualized using PyMOL software (The PyMOL Molecular Graphics System, Version 2.0 Schrödinger, LLC).

## Author contributions

SY, RO, and TT constructed the plasmids, biochemical experiments, and data analysis; TT conducted microscopic analysis and imaging; SY, MM, and TT conducted protein structure prediction in silico; MM contributed to protein structural analysis; and TT wrote the manuscript with contributions from all authors.

## Acknowledgments

We thank all members of our lab, particularly Yu Shimanuki, for their technical assistance. We are grateful to Ikuo Wada (Fukushima Medical University, Japan) for helpful discussions and providing the necessary materials. We would like to thank Editage (www.editage.jp) for English language editing. The authors declare no competing financial interests.

## Abbreviations

ATE1: arginine tRNA protein transferase 1
CHX: cycloheximide
DAPI: 4’,6-diamidino-2-phenylindole
DUB: deubiquitinase
ER: endoplasmic reticulum
ERAD: ER-associated degradation
NTAN1: N-terminal asparagine amidohydrolase 1
NTAQ1: N-terminal glutamine amidohydrolase 1
TFG: human tropomyosin-receptor kinase-fused gene
Ub: ubiquitin
WT: wild type
YFP: yellow fluorescent protein.

---

**Supplemental Figure 1.**
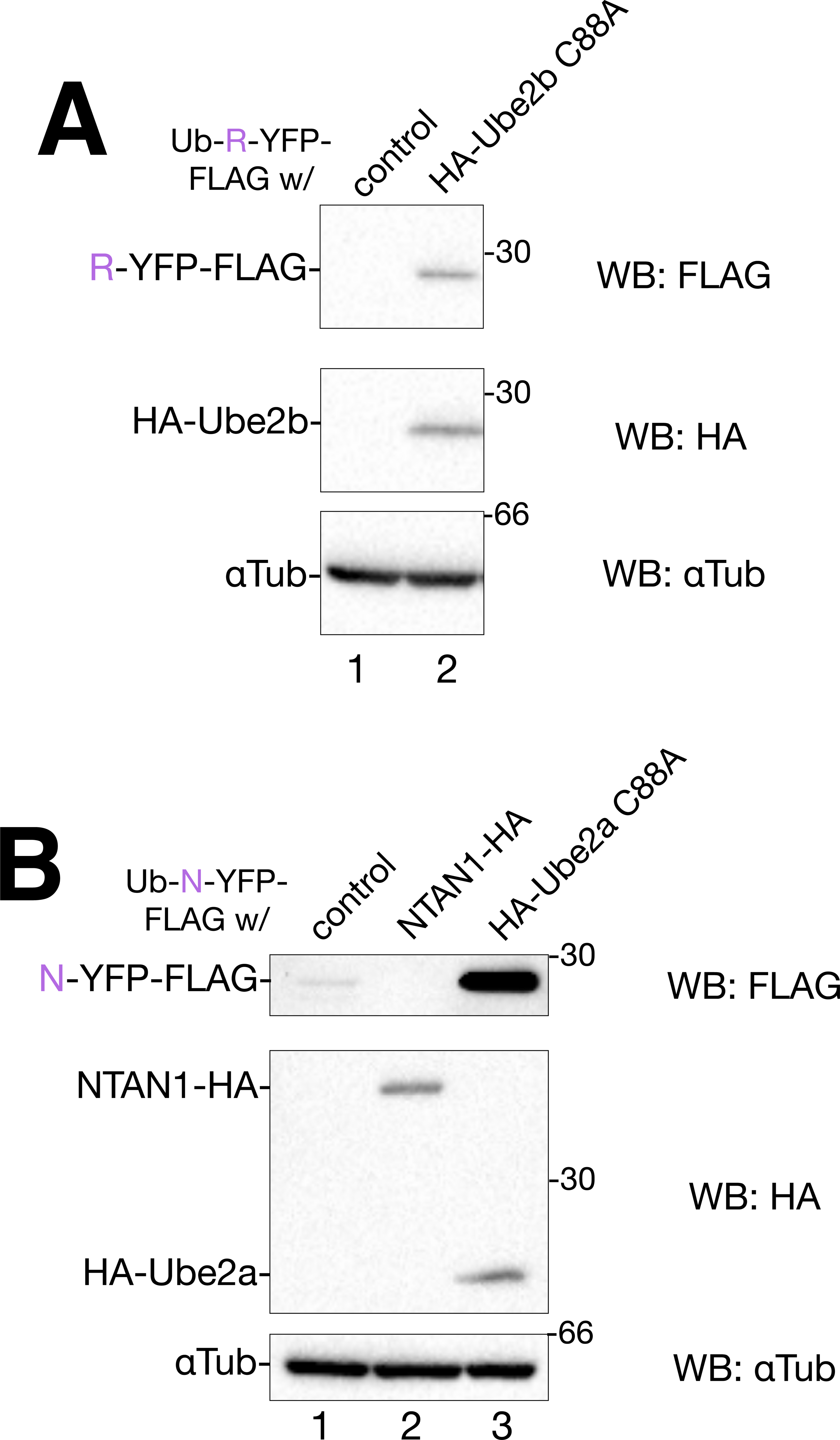
The Arg/N-end rule degradation assay in HeLa cells. (A) HeLa cells were co-transfected with Ub-R-YFP-FLAG and the control vector or HA-Ube2b C88A. The cell lysates were subjected to western blotting using the indicated antibodies. (B) HeLa cells were co-transfected with Ub-N-YFP-FLAG and the control vector, NTAN1-HA, or HA-Ube2a C88A. The cell lysates were subjected to western blotting using the indicated antibodies.

**Supplemental Figure 2.**
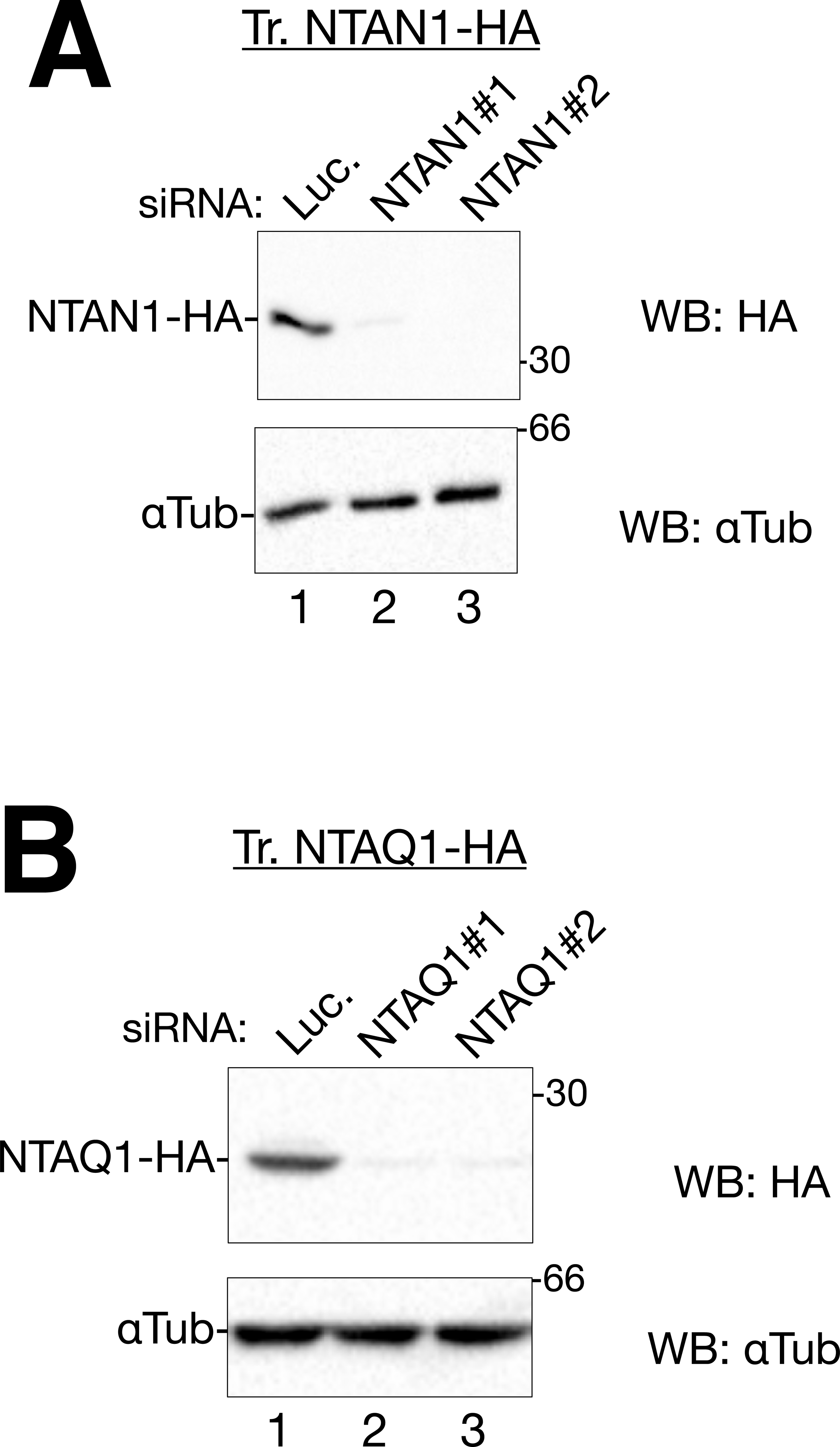
siRNA analysis for NTAN1 and NTAQ1 using exogenously expressed constructs. (A) The 293T cells were treated with a negative control (Luc.), NTAN1 #1, or NTAN1 #2 siRNA for 72 hr. Transfection of NTAN1-HA was initiated 24 h after siRNA. Immunoblotting was performed with the indicated antibodies. (B) siRNA and immunoblotting for NTAQ1. The experimental conditions were identical to those described in A.

**Supplemental Figure 3.**
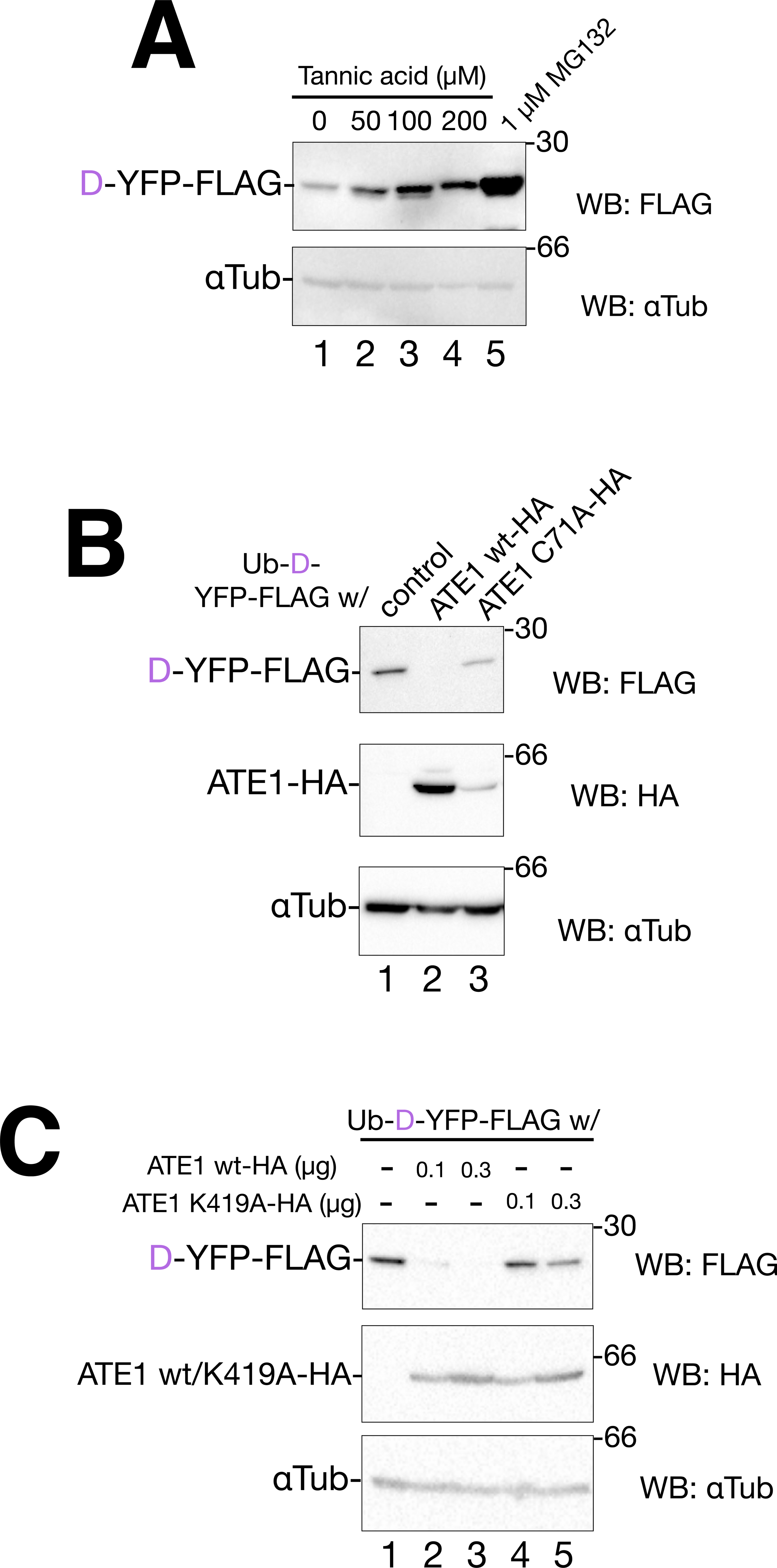
Characterization of ATE1 for D-YFP-FLAG degradation in HeLa cells by the ATE1 inhibitor tannic acid and the point mutations. (A) HeLa cells were transfected with 0.2 µg of Ub-D-YFP-FLAG with the control vector. In the last 16 h, transfected cells were incubated in the presence of tannic acid at the indicated concentration (µM, lanes 1-4) or 1 µM MG132 (lane 5). Immunoblotting was performed with the indicated antibodies. (B) HeLa cells were co-transfected with 0.2 µg of Ub-D-YFP-FLAG and the control vector (lanes 1), ATE1 wt-HA (lane 2), or ATE1 C71A-HA (lane 3), respectively. Immunoblotting was performed with the indicated antibodies. (C) HeLa cells were co-transfected with 0.2 µg of Ub-D-YFP-FLAG and the control vector (lanes 1), ATE1 wt-HA (lanes 2 and 3), or ATE1 K419A-HA (lanes 4 and 5). Immunoblotting was performed with the indicated antibodies.

**Supplemental Figure 4.**
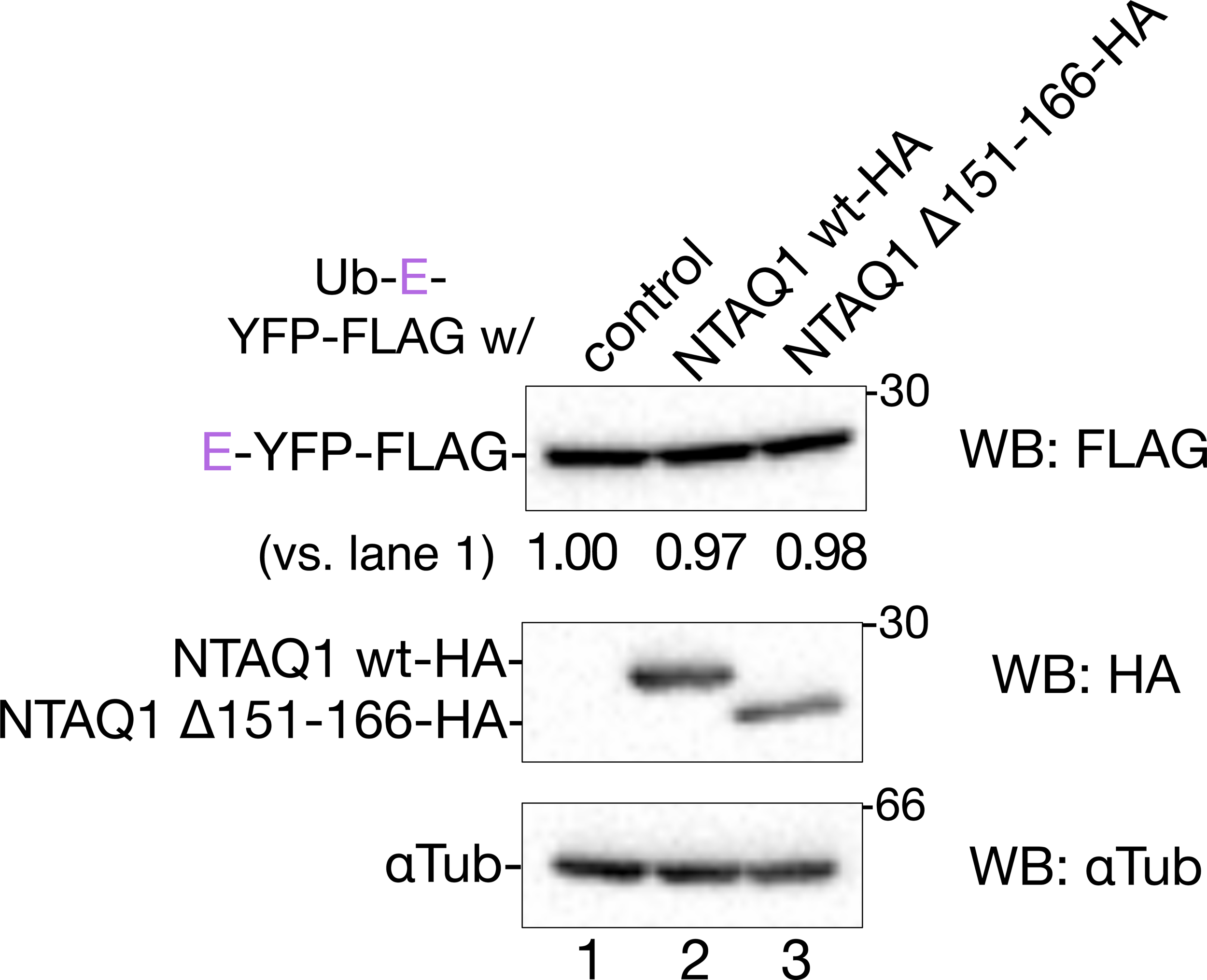
Effect of NTAQ1 on E-YFP-FLAG degradation. 293T cells were co-transfected with 0.2 µg of Ub-E-YFP-FLAG and the control vector (lanes 1), NTAQ1 wt-HA (lane 2), or NTAQ1 Δ151-166 HA (lane 3), respectively. After 48 h of transfection, cell lysates were subjected to SDS-PAGE, and the proteins of interest were detected by immunoblotting using the indicated antibodies. Values of E-YFP-FLAG are normalized by those of αTubulin and the control sample was set at 1.00.

**Supplemental Figure 5.**
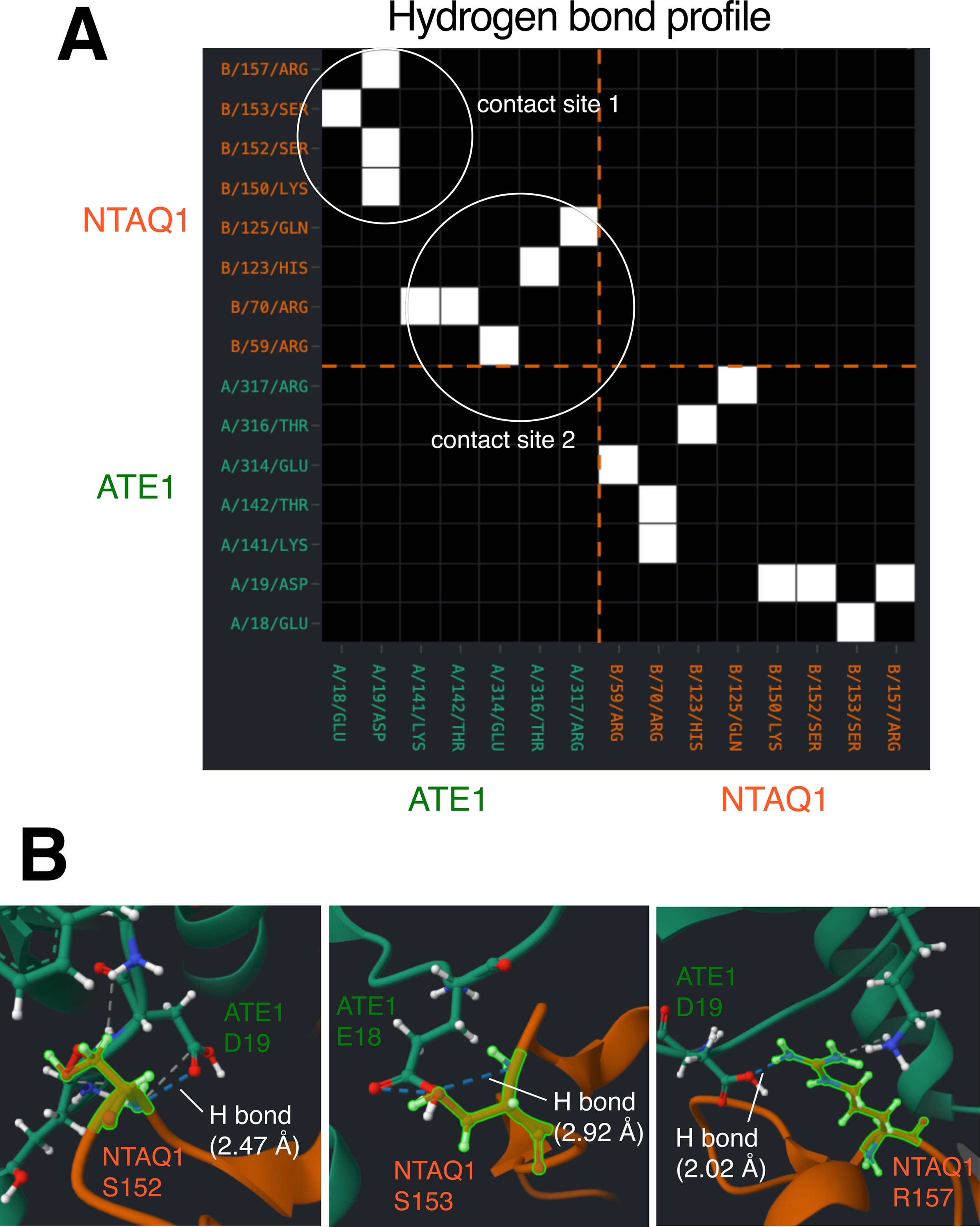
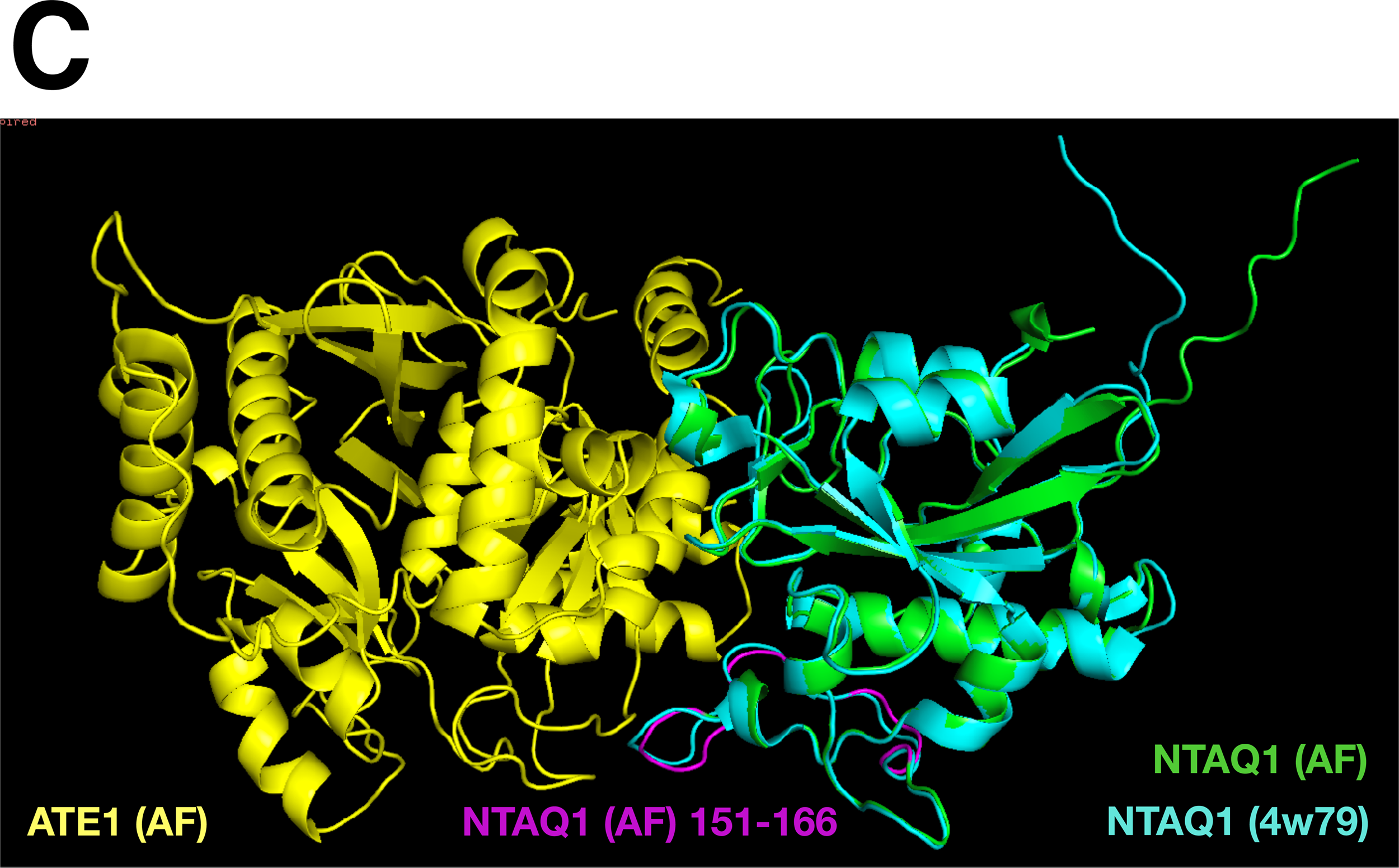
Detail analysis of ATE1-NTAQ1 complex predicted by AlphaFold2 and MMseqs2. (A) The hydrogen bond profile of the ATE1-NTAQ1 complex (Figure 8A) is shown. Amino acids contain hydrogen bonds between ATE1 (dark green) and NTAQ1 (orange) are shown in a table. Clusters of amino acids are expressed as contact sites 1 and 2. (B) Atomic contacts and hydrogen bonds (H bonds) of selected amino acids, including S152, S153, and R157 of NTAQ1, at contact site 1 are shown. The interaction at contact site 1 was visualized using the RING analysis web tool. Hydrogen bonds are indicated by a blue dashed line. (C) Merged image of the NTAQ1-ATE1 complex and NTAQ1 (PDB: 4w79). The crystal structures of human NTAQ1 and NTAQ1 in the complex generated by AlphaFold2 and MMseqs2 were aligned using PyMOL software. ATE1 by AlphaFold2 (yellow), NTAQ1 by AlphaFold2 (green), NTAQ1 from PDB 4w79 (light blue), and the loop region of NTAQ1 by AlphaFold2 (purple) are shown.

**Supplemental Table 1.**
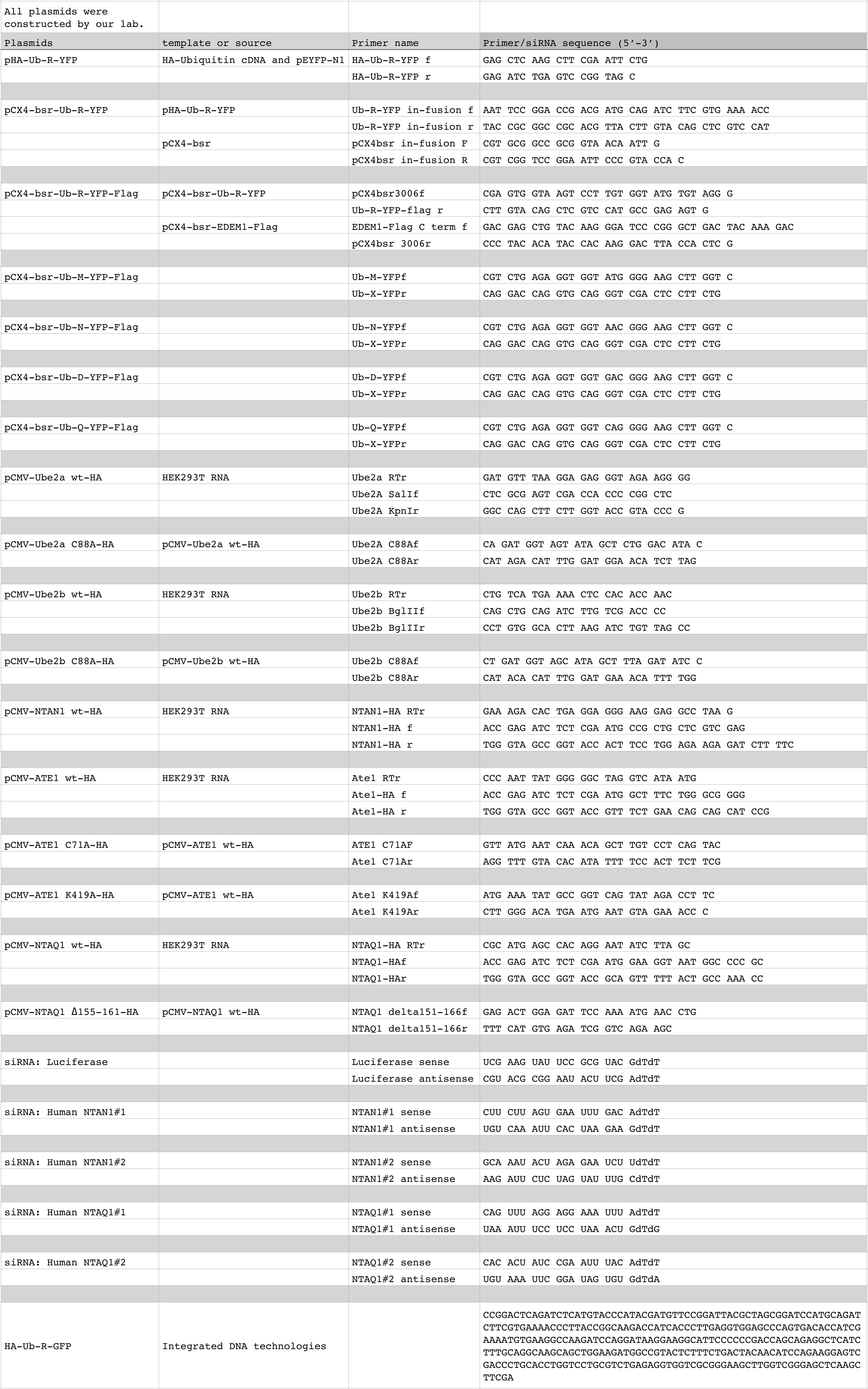

